# Metaproteomic profiling of the secretome of a granule-forming *Ca*. Accumulibacter enrichment

**DOI:** 10.1101/2024.11.06.622250

**Authors:** Berdien van Olst, Simon A. Eerden, Nella A. Eštok, Samarpita Roy, Ben Abbas, Yuemei Lin, Mark C.M. van Loosdrecht, Martin Pabst

## Abstract

Extracellular proteins are supposed to play crucial roles in the formation and structure of biofilms and aggregates. However, often little is known about these proteins, in particular for microbial communities. Here, we use two advanced metaproteomic approaches to study the extracellular proteome in a granular *Candidatus* Accumulibacter enrichment as a proxy for microbial communities that form solid microbial granules, such as used in biological wastewater treatment. Limited proteolysis of whole granules and metaproteome isolation from the culture’s supernatant successfully identified over 50% of the protein biomass to be secreted. Moreover, structural and sequence-based classification identified 387 proteins, corresponding to over 50% of the secreted biomass, with characteristics that could aid the formation of aggregates, including filamentous, beta-barrel containing, and cell surface proteins. However, while most filamentous proteins originated from *Ca*. Accumulibacter, among others cell surface proteins did not. This suggests that not only a range of different proteins, but also multiple organisms contribute to granular biofilm formation. Therefore, the obtained extracellular metaproteome data from the granular *Ca*. Accumulibacter enrichment provides a resource for exploring proteins that potentially support the formation and stability of granular biofilms, whereas the demonstrated approaches can be applied to explore biofilms of microbial communities in general.

**SIGNIFICANCE:** Biofilm-forming microbial communities are widespread and pose both challenges and opportunities in various settings in life. Structure-providing, extracellular proteins likely play a crucial role in the formation of the biofilm matrix, but these proteins are challenging to characterise due to the dynamic and complex nature of these communities. We used two advanced metaproteomic approaches to enrich for the extracellular proteins in a granule-forming *Candidatus* Accumulibacter enrichment culture as a proxy for granule-forming communities present in wastewater treatment plants. The extracellular proteins were additionally classified using structure and sequence-based annotation tools, which identified multiple different protein categories that potentially aid in granule formation, but also may provide structure to the biofilm matrix. Interestingly, although the granules were highly enriched for *Ca*. Accumulibacter, several structure-providing protein categories originated from other organisms. The obtained metaproteomic data contribute to the understanding of proteins and processes that are potentially involved in granule formation. This could further help to optimise processes involving granular biofilms and identify candidates for the recovery of novel biopolymers for biotechnological applications.

## INTRODUCTION

Microbial communities are dynamic and complex environments where individual microbes interact and compete, which can lead to synergistic effects [1, 2]. Consequently, an improved understanding of the microbial community dynamics requires a holistic and detailed knowledge of both the cellular and community functionality. This understanding will accelerate the development of strategies to control or design microbial communities to provide solutions to societal challenges [3-6].

Among the various metaomic approaches, metaproteomics is particularly powerful for community characterisation because it provides insights into the expressed proteins of different community members, their interactions, and their individual contributions to the community [7]. Characterising the complete metaproteome of microbial communities is more challenging than characterising the proteome of single, laboratory-grown cultures. For example, protein extraction can be hindered by the complex structure of the communities or the complex environment surrounding the communities [8]. Additionally, protein interference from peptides is complicated because the large number of community members results in a large and complex database [9, 10]. Furthermore, metagenomic approaches for constructing comprehensive reference sequence databases are highly dependent on the chosen pipeline, and their completeness is difficult to verify [9, 11, 12].

Characterising microbial community proteins beyond those involved in central metabolic pathways presents an additional challenge, particularly when analysing structure-providing proteins. Such proteins can have unconventional properties or modifications that challenge their detection [13] and their functions are difficult to verify in expression studies resulting in many unknown members. However, structure-providing proteins are crucial in many microbial communities. For example, the crosslinking ability of proteins and other biopolymers aids in the formation of an extracellular matrix that aggregates the community together in a biofilm [1, 14, 15]. The biofilm facilitates intercellular interaction, enhances water retention and nutrient sorption, and protects the microbial community against various environmental stresses [1, 2]. On the other hand, biofilm-forming communities can pose challenges and opportunities in among others medical and industrial settings [16, 17]. Therefore, it is important to understand the role of proteins beyond the central metabolic pathways.

The proteins in the extracellular matrix may play a crucial role in fine-tuning microbial communities in both natural and engineered ecosystems. For example, in biological wastewater treatment, water is cleaned from pollutants such as carbon, phosphorus and nitrogen by microbial communities that are separated from the cleaned water by gravitational settling. These communities first form flocs that only settle slowly but can be transformed into dense granules that settle much faster, thereby offering various operational and economic advantages for wastewater treatment plants [18-20]. Interestingly, these granules were reported to have a higher protein content than flocs with the centre of the granules mainly consisting of proteins [21]. Furthermore, while certain exopolysaccharide components were found to be important for granule structure in various granule types [22, 23], this specific component was absent in granules enriched in *Ca*. Accumulibacter [23]. This might indicate that other components, such as proteins, play an important role in the formation of these granules. *Ca*. Accumulibacter has been identified as the dominant organism in dense granular wastewater treatment plant communities that perform enhanced biological phosphorus removal [11, 24]. Although this organism cannot be isolated from the community, it can be enriched by selective growth conditions [25], thereby reducing the complexity of the proteomic pool. Therefore, granular Accumulibacter enrichment cultures serve as valuable models for studying extracellular proteins and their potential roles in biofilm formation.

The potential processes involved in granule formation in Accumulibacter enrichment cultures have been studied previously. For instance, Barr and colleagues examined the metaproteomic changes occurring during the transition from microbial flocs to granules and identified proteins related to stress and to outer membrane proteins that allow scavenging for extracellular nutrients [26]. Another study measured the turnover of extracellular proteins and polysaccharides by tracking the incorporation of stable isotopes and found that the extracellular biopolymer turnover was comparable to the intracellular biopolymer turnover [27]. However, the harsh extraction procedures resulted in the release of intracellular proteins [28] and only 20% of the total protein intensity was regarded as extracellular [27].

Several other studies that investigated the extracellular matrix also observed large fractions of intracellular proteins [29-33]. Therefore, a gentler approach is needed to selectively measure the proteins in the extracellular space and exposed at the surfaces of the granules.

To more selectively measure the extracellular proteome of a granular *Ca*. Accumulibacter enrichment culture, we employed two metaproteomic approaches that target the extracellular space, with a particular focus on identifying proteins that contribute to structural features. The first approach focused on the supernatant of the enrichment to study proteins that can also enter the intergranular space. The supernatant has been previously employed to study soluble extracellular proteins, albeit requiring large sample volumes and prefractionation with SDS-PAGE before MALDI-TOF identification of protein bands [31]. The second approach employed in our study utilized limited proteolysis of whole granules, allowing for the cleavage of fragments from proteins that are exposed to the extracellular space [34]. The metaproteomic data were analysed using a metagenomics constructed database from the same enrichment and were classified with a range of protein function and structure prediction tools. Together, these two approaches reveal a large spectrum of proteins that are released into the extracellular space, of which some may support aggregation and the development into a granular biofilm.

## METHODS

### Reactor Operation

The Accumulibacter enrichment culture was cultivated in a 1L laboratory-scale sequencing batch reactor controlled by a Applikon System 10 controller. The inoculum originated from the return flow (return activated sludge) of the Harnaschpolder wastewater treatment plant in Den Hoorn, The Netherlands. The sequencing batch reactor operated in six-hour cycles. Each cycle started with a 10 min nitrogen sparging phase to remove any dissolved oxygen from the liquid, followed by a 30 min feeding phase, a 105 min anaerobic phase, and a 135 min aerobic phase. In the last minute of the aerobic phase, approximately 30 mL of sludge was discharged, followed by a 30 min settling phase and a 50 min effluent discharge phase, after which the cycle was repeated. The reactor was stirred at 600 RPM in all phases, except for the settling and the effluent discharge phases. The culture was fed with 50 mL COD medium (124 mM Sodium Acetate), 50 mL mineral medium (20 mM Ammonium Chloride, 12 mM Magnesium Sulphate, 5 mM Calcium Chloride, 13 mM Kalium Chloride, 13 mM Sodium Dihydrogen Phosphate, 0.69 mM N-Allylthiourea (C_4_H_8_N_2_S), 18 mL of Trace elements [35] and 500 mL of distilled water with Bacto™ yeast extract (10 mg/L). The pH was set to 7.2 (±0.05), maintained by addition of 0.5 M HCl (with antifoam) and 0.5 M NaOH.

### Sampling

Samples were taken in the last minutes of the aerobic phase, while the reactor was still stirred. A large sample was drawn, and the granules were separated from the supernatant by settling for a minute. The granules for whole granule proteome and metagenome analysis were immediately frozen and stored at -20 °C until further processing, whereas the granules for shaving were freshly processed (see below). The supernatant (1.5 mL) was centrifuged (14.000 rcf, 3 min, 4 °C) and the proteins were precipitated in triplicate with trichloroacetic acid (TCA; 4:1 ratio SN:TCA) at 4 °C for 30 minutes, pelleted by centrifugation (14.000 rcf, 15 min, 4 °C), and subsequently stored at -20 °C until further processing. In addition to the aerobic samples, metagenome and supernatant samples were taken in the last minutes of the anaerobic phase in the same cycle and processed analogous to the aerobic samples.

### Limited proteolysis of granules (“enzymatic granule shaving”)

Granules were shaved by either Trypsin, Chymotrypsin, LysC or LysC/Trypsin in triplicate. Fresh granules (30 mg) were incubated with 15 μL of 0.1 μg/μl protease in 500 μL 100 mM Ammonium Bicarbonate (ABC) in a ThermotopMixer with Thermotop (37 °C, 300 rpm). The LysC/Trypsin samples were initially digested with LysC, and Trypsin (5 μL of 0.1 μg/μl) was added after two hours to prevent digestion of LysC by Trypsin. Granule damage was checked every hour by comparing to a control incubation without protease. After four hours of incubation, the granules were collected (2 min, 5000 rcf) and the supernatant was transferred to clean LoBind Eppendorf tubes. To promote denaturation of the shaved-off proteins, 180 mg urea (corresponding to 6 M) was dissolved in the supernatant of the LysC samples. After an overnight incubation, the samples were centrifuged (2 min, 14000 rcf) to ensure no cell debris was transferred to the downstream processing. The digested sample was cleaned by solid phase extraction (SPE) on an Oasis HLB 96-well μElution Plate (Waters). The wells were conditioned in 100% MeOH and equilibrated with pure LC-MS H_2_O. Following equilibration, the samples were loaded onto the columns, washed twice with 5% MeOH in LC-MS H_2_O and eluted with 2% formic acid in 80% MeOH and subsequently 1 mM ABC in 80% MeOH. The collected flow-through was transferred to LoBind Eppendorf tubes and dried in a centrifuge concentrator at 45 °C. The dried sample was dissolved in 3% acetonitrile in 0.1% formic acid in LC-MS H_2_O to a concentration of approximately 0.25 mg/mL as estimated from the absorbance at 280 nm using a Nanodrop.

### Protein extraction from whole granule samples

Granules for whole granule proteome analysis were lysed in duplicate by combining 40 mg granules, 150 mg acid-washed glass beads, 175 µL of B-PER reagent, and 175 µL of 1% NaDOC in 50 mM TEAB pH 8 followed by three rounds of bead beating for 90 seconds with intermediate cooling on ice for 30 seconds. The cell suspension was subsequently incubated for 3 minutes at 80 °C in a ThermotopMixer with Thermotop (1000 rpm) followed by 10 minutes of sonication in an ultrasonic bath. The remaining cell debris was pelleted (10 min, 14000 rcf, 4 °C), the supernatant was transferred to a clean LoBind Eppendorf tube, and the proteins were precipitated with TCA (ratio 4:1) at 4 °C for 30 minutes followed by centrifugation (14.000 rcf, 15 min, 4 °C).

### Sample processing for whole granules and supernatant samples

Both the whole granule and the supernatant protein pellets were washed with 200 µL ice-cold acetone (3 min, 14000 rcf) and dissolved in 6 M Urea (granules: 100 µL; supernatant: 50 µL). The dissolved pellet was reduced with 10 mM dithiothreitol (DTT; granules: 30 µL; supernatant: 5.5 µL) for 60 minutes using a ThermotopMixer with Thermotop (37°C, 300 rpm) and alkylated with 20 mM Iodoacetamide (IAA; granules: 30µL; supernatant: 5.5 μL) for 30 minutes in the dark. Subsequently, the samples were diluted in 100 mM ABC (granules: 285 µL; supernatant 440 µL) digested with either 0.1 μg/μl trypsin or 0.1 μg/μl chymotrypsin (granules: 15 µL; supernatant: 10 µL) for approximately 18 hours in a Thermomixer with Thermotop (37°C, 300 rpm). The digested sample was cleaned by SPE as described for granule shaving and concentrated to approximately 0.5 mg/mL and 0.25 mg/mL for granules and supernatant samples, respectively. The whole granule samples digested by trypsin were injected twice in the LC-MSMS.

### Metaproteomics

Metaproteome samples were loaded on an EASY nano-LC 1200 equipped with an Acclaim PepMap RSLC RP C18 separation column (50 μm × 150 mm, 2 μm and 100 Å). For the granule-based samples, the flow rate was maintained at 350 nL/min with 0.1% formic acid in water (solvent A) and a 120 min linear gradient from 5 to 65 of 80% acetonitrile and 1% formic acid in water (solvent B). The elute was sprayed Nanospray Flex (Proxeon) in a Thermo Q Exactive Plus mass spectrometer (Thermo Scientific, Germany) operated in data dependent acquisition (DDA) mode. Full spectra were recorded with peptide signals between 385 and 1250 m/z at a 70K resolution with an automatic gain control of 3e6 and a maximum injection time of 75 ms. The top 10 most abundant precursor ions with a charge state between 2-5 were fragmented (28 % normalised collision energy, 2.5 m/z isolation window, and 0.2 m/z isolation offset) and recorded at a 17.5K resolution with an automatic gain control of 2e5 and a maximum injection time of 100 ms. The supernatant samples were analysed with a 60 min linear gradient from 2 to 40% of solvent B and full spectra were recorded between 350 and 1250 m/z with a maximum injection time of 100 ms. The top 10 most abundant precursor ions with a charge state between 2 and 5 were fragmented (3.0 m/z isolation window) with an automatic gain control of 5e5. The mass spectrometric raw data was analysed per approach using PEAKS Studio (Bioinformatics Solutions Inc., Canada) and searched against the metagenomic-constructed database (see below) and the cRAP contaminant database. A maximum of two missed cleavages was allowed for the whole granule and supernatant samples and four missed cleavages for the shaved samples. The digestion mode was set at specific for the whole granule and supernatant samples, and semi-specific for the shaved samples. Carbamidomethylation (C) was set as fixed modification on samples that were alkylated (i.e., whole granule and supernatant samples), whereas deamidation (NQ), oxidation (M) and acetylation (Protein N-term) were set as variable modification in all samples. Peptide identifications were filtered for 1% false discovery rate, protein identifications required at least one unique peptide and the top proteins per protein group were exported. The identified proteins were grouped across the different approaches if they were assigned to the same protein group in at least one approach and were not assigned to different protein groups in any of the approaches. The metaproteome intensities were converted to iBAQ using the theoretical number of peptides per protease [36].

### DNA extraction from granules

DNA extractions were performed in duplicate based on the Phenol/Chloroform extraction as described by Trigodet et al. [37]. Briefly, 80 mg of frozen granules was homogenised using a potter device in a 2 ml Eppendorf and incubated at 37 °C for 1 hour in 1 ml TLB (10 mM Tris-HCL pH 8.0, 25 mM EDTA pH 8.0, 100 mM NaCl, 0.5% [w/v] SDS, 20 μg/ml RNase A). Subsequently, the samples were incubated with Proteinase K (final concentration 200 µg/ml) at 50 °C for 2 hours with gentle inversing every 30 minutes. The lysate was transferred to a 2 ml phase-lock tube (Quantabio, Germantown MD, USA) containing 800 µL Phenol, inverted for 10 minutes and centrifuged for 15 min at 17000*g, which was repeated with 800 µL Phenol/Chloroform (PCl 25:24:1). The DNA was precipitated by adding 720 µL (0.8 vol) 100% isopropanol and centrifuging for 15 min. The resulting pellet was washed twice with 1 mL 70% ethanol and resuspended overnight in TE-buffer. The resuspended sample was cleaned with a Qiagen Genomic Tip-20 (Hilden, Germany) according to the manufacturer’s protocol with isopropanol precipitation. The high molecular weight fraction was subsequently selected based on the method described by Stortchevoi et al. [38]. In short, 4000 ng DNA was incubated with 10 mM TRIS pH8, 0.14 vol Mg-PEG buffer (70 mM MgCl_2_ in 0.7% (v/v) Polyethylene glycol 8000) and washed AMPure XP beads (Beckman Coulter) for 60 minutes while shaking to keep the beads in suspension. The beads were collected on a magnet and washed twice with 500 µL 80% Ethanol. The DNA was released from the beads by incubating the beads in 50 µL H_2_O for 10 min at 37 °C while shaking (500 rpm) and stored at -20 °C until further processing. Part of the DNA was loaded onto an agarose gel and electrophoresis was performed to evaluate the size of the DNA and its suitability for Illumina library preparation.

### Metagenomics

DNA libraries were prepared using the Nextera XT DNA library preparation kit (Illumina) and sequenced on an Illumina NovaSeq6000 with a paired-end 150bp strategy (Baseclear, Leiden, The Netherlands) for 10GB. The data quality was assessed, the metagenome was assembled using MEGAHIT (v 1.2.9) [39] and genes were predicted and annotated using Prokka (v 1.14.5). The results obtained by the four samples were clustered to reduce sequence redundancy using CD-HIT [40] at 95% sequence identity level, resulting in a single database that was used for protein identification.

### Processing of metaproteomic data

The identified proteins were annotated with information about their location, taxonomy, functionality and sequence properties. Secreted proteins were identified by predicting the presence of a signal peptide using SignalP (v 6.0) [41], whereas transmembrane domains were predicted using DeepTMHMM (v 1.0) [42]. The location of the identified proteins was further predicted by PSORTb (v 3.0.3) [43] and by DeepLocPro (v 1.0) [44], a prokaryotic extension to DeepLoc [45]. The taxonomic origin was predicted using GhostKoala [46] and further validated for the identified aggregating proteins by DIAMOND [47]. Proteins were functionally annotated using Eggnog mapper (v 2.1.12) [47-49]. Sequence-based protein properties were calculated using the ProtParam module in the SeqUtils package of Biopython (v 1.78), whereas protein structural properties were calculated using NetSurfP (v 3.0) [50]. To classify proteins as potentially aggregating, we ranked all measured proteins by the properties describing charge, hydrophobicity (GRAVY), secondary (percentage of sequence in beta-sheet) and tertiary (percentage of cysteines) structures, aromaticity, protein length, surface accessibility and disorder. Subsequently, proteins were selected as aggregating if (i) at least one of these properties belonged to the top ten percent (GRAVY, beta-sheet percentage, cysteine percentage, aromaticity, protein length), or bottom ten percent (surface accessibility and disorder), (ii) the protein abundance belonged to the top fifty percent in at least one approach and (iii) the protein was predicted to be secreted or extracellular. The selected proteins were clustered using a Euclidean distance and a complete clustering method. As both a negative and a positive charge might contribute to aggregating, the protein charges were converted to absolute numbers before selecting the top ten percent, but the non-converted values were used for clustering.

## DATA AVAILABILITY

The metaproteomic raw data have been deposited to the ProteomeXchange Consortium via the PRIDE partner repository with the dataset identifier PXD057220.

## RESULTS

Here we employed two different metaproteomic approaches to identify and characterise the secretome of a *Ca*. Accumulibacter enrichment (hereafter Accumulibacter) as a proxy for aerobic granular sludge employed in wastewater treatment, with a focus on identifying potential aggregate forming proteins that play a role in the formation and structure of the granular matrix. As Accumulibacter cannot be cultured in pure form [25], we enriched a sample obtained from a wastewater treatment plant for Accumulibacter by cyclic anaerobic feeding of acetate and aerobic growth phases in a bioreactor. Subsequently, the enrichment was analysed by metagenomics and metaproteomics, where the metagenome served as a reference for database searching experiments. Two metaproteomic approaches were employed to identify the secreted metaproteome of the enrichment cultures, namely “enzymatic granular shaving” and “supernatant proteomics” (Figure 1A). The first approach aimed to identify proteins inside the granule’s biofilm by directly adding proteases to intact granules, thereby ‘shaving’ the granules of their extracellular and transmembrane proteins. This approach relies on the digestion of native proteins, which are not easily accessible for proteases. Therefore, multiple proteases were employed in parallel to enhance protein identification (Supplemental Figure 2). The second approach aimed to identify proteins in the culture’s supernatant, which is expected to be enriched in secreted proteins. For this supernatant approach, we compared samples harvested from the anaerobic feeding phase and the aerobic growing phase, which showed only minimal proteome differences between these two phases (Supplemental Figure 3). These two approaches were compared to the proteome obtained from lysed whole granules (Supplemental Figure 1).

**Figure 1:**
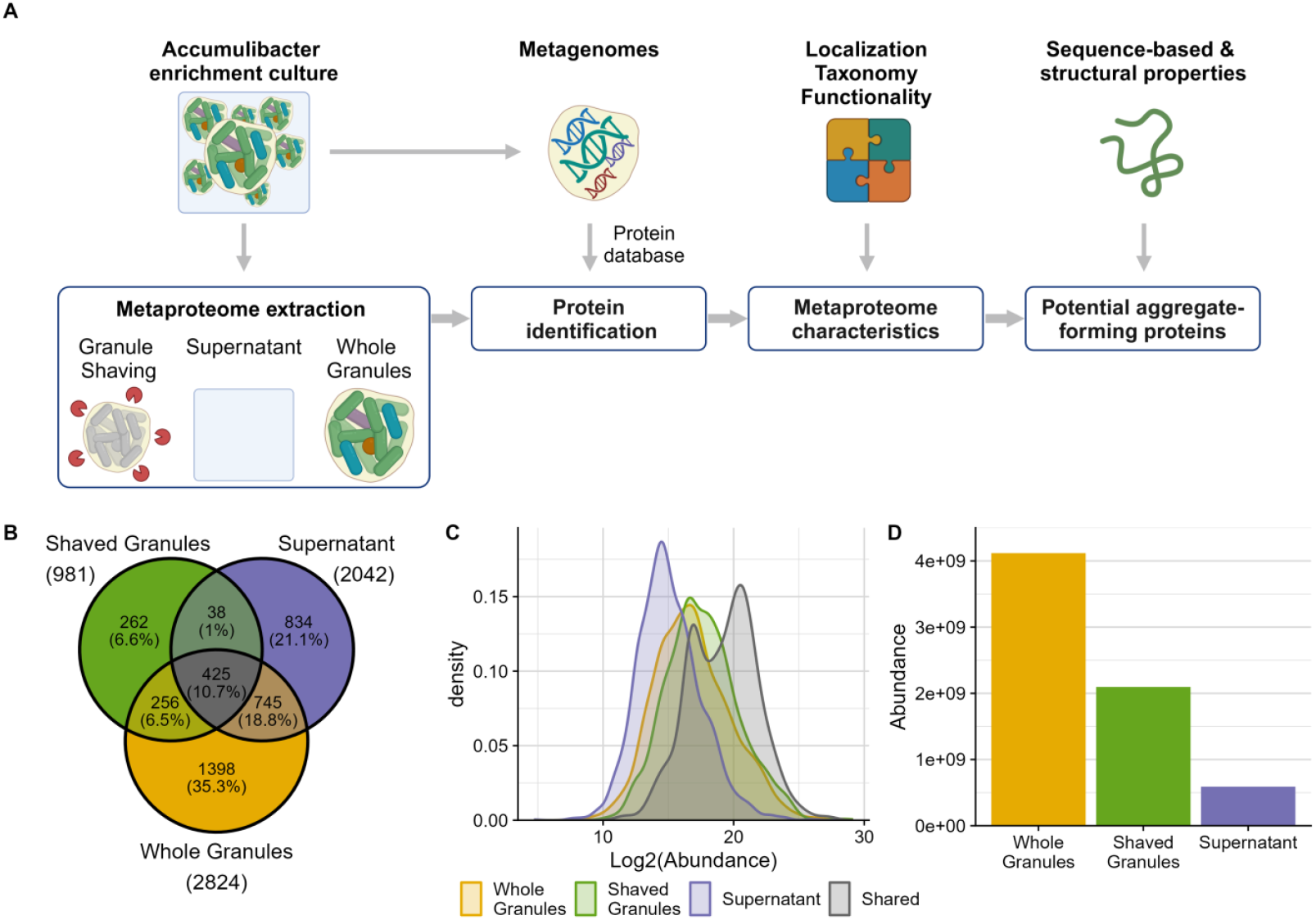
Workflow and global comparison of different metaproteomics approaches applied to Ca. Accumulibacter enrichment. (A) Workflow (B) Venn diagram displaying the identified protein groups per approach and the overlap between them. (C) Overlap in the log2 protein abundance (x-axis) for the protein groups identified by the different approaches and the protein groups shared by all approaches (colours). (D) Total protein biomass calculated from the summed protein abundances (y-axis) per approach (x-axis).

### The extracellular metaproteome of a *Ca*. Accumulibacter enrichment

The three different approaches identified in total almost 4000 protein groups, of which 71% was identified in the whole granules, 52% was identified in the supernatant, 25% was identified in the shaved granules, and 11% was identified in all approaches (Figure 1B). These shared protein groups were mostly high-abundant but also included some protein groups with average abundances (Figure 1C). Only 1% of the identified protein groups were found in both the shaved granules and the supernatant, which correlated only moderately (Pearson: 0.41; Supplemental Figure 4A), and both approaches shared more proteins and correlated better with the whole granule approach (Pearson: 0.58 for shaved granules and 0.57 for supernatant; Supplemental Figure 4A). Each approach also identified a unique set of protein groups (Figure 1B) which were mainly average and low abundant (Supplemental Figure 4B). Notably, the protein biomass (i.e., summed total protein abundance) was two times lower in the shaved granule samples compared to whole granule samples (Figure 2D), which corresponds to the 2-fold lower amount of proteolytic digest analysed. Conversely, the protein biomass observed in the supernatant samples was 7-fold lower compared to the whole granules, which likely reflects both the lower injection volume and the shorter gradient used to analyse the samples, thereby providing fewer identifications.

**Figure 2:**
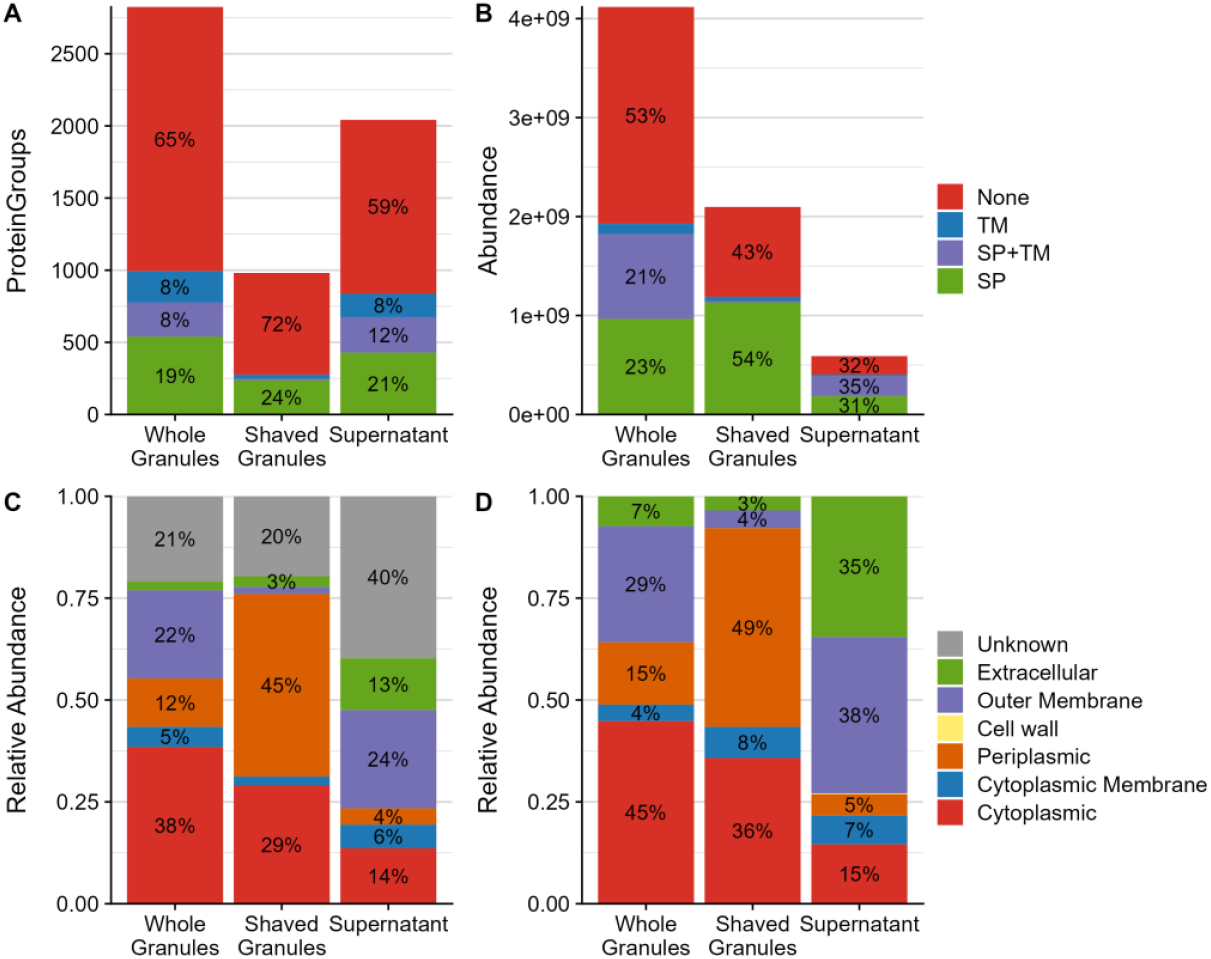
Enrichment of secreted and extracellular proteins by the different metaproteomics approaches. (A) The number of identified protein groups (y-axis) and (B) the summed protein abundance (y-axis) of proteins with a predicted signal peptide (SP; green), a predicted transmembrane domain (TM; blue), both predictions (SP+TM; purple) or no predictions (None; red) for the different approaches (x-axis). (C, D) The relative abundance of proteins (y-axis) predicted at different cellular locations (colours) as predicted by (C) PSORTb and (D) DeepLocPro for the different approaches (x-axis).

The shaving and supernatant approaches were developed to investigate the proteins that are secreted by the microbes in the enrichment. Therefore, we examined the proteins identified in each approach for the presence of transmembrane domains and signal peptides that indicate secretion. These tools predicted that of the almost 4000 protein groups identified by all approaches, 19% had a signal peptide, 8% had a transmembrane domain and 8% had both. These percentages were similar for the whole granule and supernatant approaches (Figure 2A), indicating that a substantial fraction of the identified protein groups can be regarded as intracellular. Conversely, 25% of the protein groups in the shaved granules were predicted to be secreted and only a limited number of transmembrane proteins was identified. The lack of transmembrane proteins in the shaved granules also partly explains the substantial number of proteins that was only identified in whole granules and the supernatant (Figure 1B) of which 30% was predicted to have a transmembrane domain. Interestingly, the small number of secreted proteins accounted for the majority of the protein biomass. Of the shaved biomass, 54% was predicted to be secreted and transmembrane proteins contributed only marginally (Figure 2B) while 66% of the supernatant biomass was predicted to be secreted, of which almost half was also predicted to have a transmembrane domain. In the whole granule samples, only 43% of the protein biomass was predicted to be secreted, of which half was also predicted to have a transmembrane domain, similar to the supernatant samples. Moreover, in the shaved granules, 50% and 75% of the secreted protein biomass was covered by 2 and 9 proteins, respectively. Conversely, in the supernatant, 6 and 26 proteins covered over 50% and 75% of the secreted protein biomass, respectively, which is similar to the whole granules where 4 and 21 proteins covered 50% and 75% of the secreted protein biomass, respectively.

The absence of a predicted secretion signal does not necessarily mean that the proteins are not secreted, as there might be alternative routes to the extracellular space. Therefore, the protein location was further examined with PSORTb and DeepLocPro (a prokaryotic extension to DeepLoc). Both these tools confirm the high number but low abundance of cytoplasmic protein groups in shaved granules and the supernatant, albeit with different percentages (Figure 2C and D; Supplemental Figure 5). Moreover, they validate the substantial non-cytoplasmic fraction in whole granules. Interestingly, both tools indicate that the different approaches enrich for proteins at different locations (Figure 2C and D). Most biomass in the shaved granules is assigned to periplasmic protein groups, whereas most biomass in the supernatant is assigned to outer membrane proteins. In addition, a large fraction of the supernatant biomass is assigned to extracellular proteins when predicting with DeepLocPro, which is substantially less when predicting with PSORTb. In whole granules, the biomass mainly contains cytoplasmic proteins, followed by outer membrane and periplasmic proteins. Hence, while the shaved granule and supernatant approaches still result in a significant number of intracellular proteins, these proteins only account for a small fraction of the protein biomass confirming that both approaches enrich for secreted proteins, although from different cellular locations.

The secreted proteins are investigated in an Accumulibacter enrichment culture, as Accumulibacter is expected to be involved in dense granule formation [11, 24]. To examine the Accumulibacter enrichment, we investigated the culture’s taxonomy as identified by the different approaches. The two approaches that immediately target granules displayed similar enrichment patterns across the different taxonomy levels and indicated a high enrichment (96%) of the Pseudomonodota phylum, the Betaproteobacteria class (∼89%) and the *Ca*. Accumulibacter genus (∼78%; Figure 3). Conversely, only 88% of the protein biomass in the supernatant belonged to the Pseudomonodota phylum, only 66% to the Betaproteobacteria class and only 43% to the *Ca*. Accumulibacter genus. The remaining proteins in the supernatant could be attributed to more than 100 different taxonomies (based on protein sequence alignment) but were mostly low abundant (< 2%). Nevertheless, 2% of the protein biomass could be attributed to viruses, 6% to the Bacteroidota phylum, 17% to the Gammaproteobacteria class, and 5% to the Hydrogenophaga genus, suggesting the presence of planktonic cells in the bioreactor that are not directly part of the granular community, and such cells were indeed visible in the bioreactor culture (data not shown). Hence, while the granule community is highly enriched for Accumulibacter, this was not reflected in the identified supernatant proteome.

**Figure 3:**
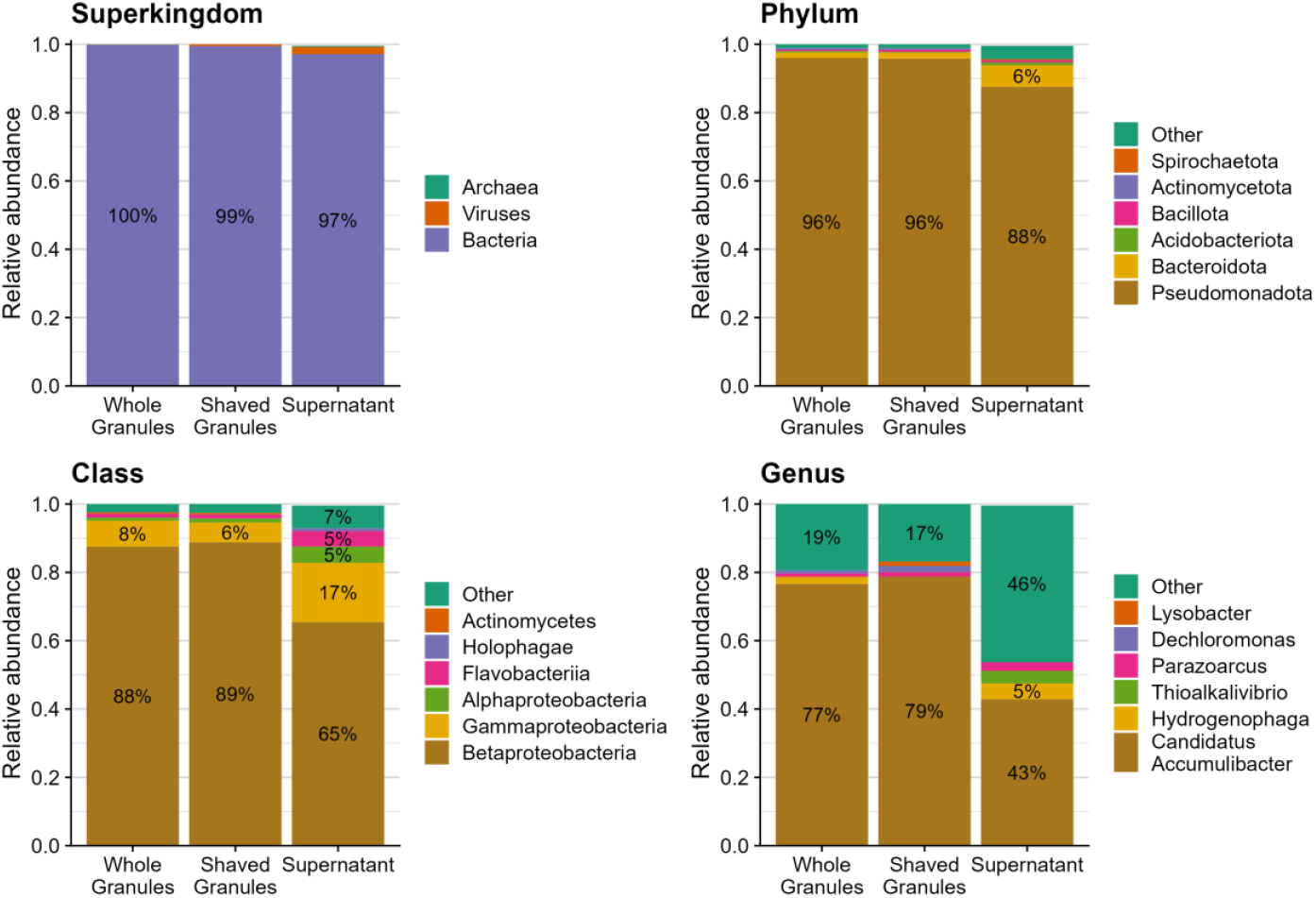
Taxonomic composition obtained from different metaproteomics approaches for Ca. Accumulibacter enrichment. Enrichment for different organisms based on relative contribution to the protein biomass (y-axis) in the different approaches (x-axis) on different taxonomy levels (panels).

### Functional classification of identified proteins

To gain more insight in the protein functions, proteins were further assigned to Clusters of Orthologous Groups (COGs) [51, 52]. Of the 4000 identified protein groups, 21% could not be assigned to a COG group and 12% had an unknown function (S category). The missing functions predominantly impacted the supernatant samples as 45% of the protein biomass and 51% of the secreted and extracellular protein biomass could not be assigned (i.e., either NA or S; Figure 4). The most abundant protein group (10%) without assignment corresponded with a hypothetical protein from Accumulibacter with a secretion signal, which was also found in whole granules (2%) but not in shaved granules (Supplemental Table 1). The most abundant group (3%) with unknown function corresponded to a tail sheath protein that was only found in the supernatant, which was predicted to originate from Hydrogenophaga but also shared a 100% sequence identity with a major tail sheath protein from Accumulibacter. The “Cell wall, membrane and envelop biogenesis” category (M) was the most abundant assigned category in the whole granules and supernatant, covering about 21% of the protein biomass, which corresponded to 44% and 28% of the secreted and extracellular protein biomass, respectively (Figure 4). In contrast, only 6% of the secreted and extracellular protein biomass in the shaved granules belonged to this category, which was explained by the high number of transmembrane proteins that were only limitedly identified in the shaved granules. The main contributing proteins in this category were various high-abundant, secreted, outer membrane porins (Supplemental Table) that cover 19% and 17% of the protein biomass in the whole granules and supernatant, respectively, making it the most abundant proteins in these approaches. The “Energy production and conversion” category (C) was the most abundant fraction in the shaved granules, covering 31% of the protein biomass and 46% of the secreted and extracellular protein biomass, with the latter being almost 14 and 44 times higher compared to the whole granules and supernatant, respectively. The main contributing protein in this category was a secreted cytochrome C from Accumulibacter, which was also the most abundant protein in the shaved granules covering 23% of the protein biomass. The “Signal transduction mechanisms” category (T) also covered a large fraction of the secreted and extracellular protein biomass in both the shaved granules (24%) and whole granules (11%), whereas it only covered 2% in the supernatant. The main contributing protein in this category was a secreted, hypothetical protein with a Calcium/Chemotaxis domain. This protein was the second and third most abundant protein in the shaved and whole granules, making up 12% and 4%, respectively. Both the “Cell motility” and “Intracellular trafficking, secretion, vesicular transport” categories (N and U) covered ∼12% of the secreted and extracellular protein biomass in the supernatant, which was over 2-fold and 9-fold larger than in the whole and shaved granules, respectively. The main contributing proteins in these categories were various fimbrial proteins that are involved in cell adhesion and assigned to both classes. Interestingly, the most abundant fimbrial protein was assigned to Accumulibacter, which was twice as abundant as the fimbrial proteins of two other Betaproteobacteria. Taken together, the approaches enriched for different proteins in different functional categories although there was some overlap.

**Figure 4:**
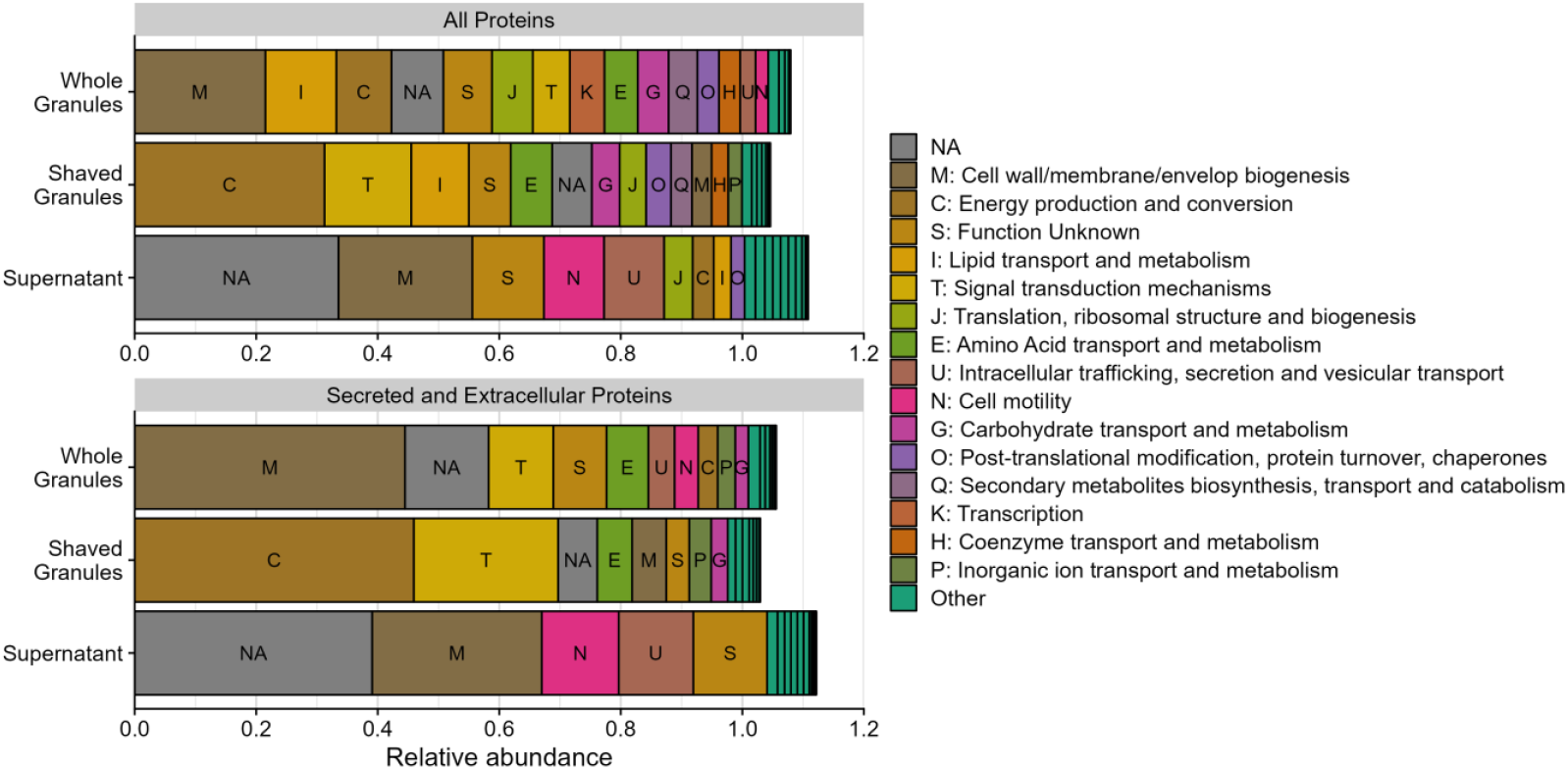
Relative protein biomass of Clusters of Orthologous Groups (COGs) for whole granule, shaved granule, and supernatant metaproteomes. Distribution of the relative protein abundance (x-axis) per metaproteomic method (y-axis;) over different COG categories. The upper panel shows this for the total protein biomass and the lower panel for the protein biomass predicted to be secreted or extracellularly located. Categories with a relative abundance lower than 2% are coloured as “Other”. Proteins annotated with more than one COG category are allocated to all those categories, resulting in a total relative abundance larger than 1.

### Large spectrum of proteins with aggregate-forming properties

To gain more insight in proteins that form and contribute to the structure of the granules, the secretome was investigated for proteins with properties that allow aggregation or non-covalent interaction forces that can keep a biofilm matrix together [14]. We defined aggregating properties as the intrinsic sequence-based properties that contribute to the formation of secondary, tertiary or quaternary structures and therefore allow self-association or subunit-association. To this end, we predicted the net charge, hydrophobicity (GRAVY), aromaticity, cysteine percentage and protein length and predicted the beta-sheet percentage, relative surface accessibility and disorder. Proteins were considered to be potentially aggregating when at least one of the properties was in the top ten percent of the measured proteins and the protein abundance was in the top fifty percent in at least one approach. To further narrow down our search, we only selected proteins that were predicted to be secreted or extracellular as granule formation is expected to be intentional. This resulted in a list of 387 protein groups that were clustered together into 9 main clusters (Figure 5; Supplemental Table 2).

**Figure 5:**
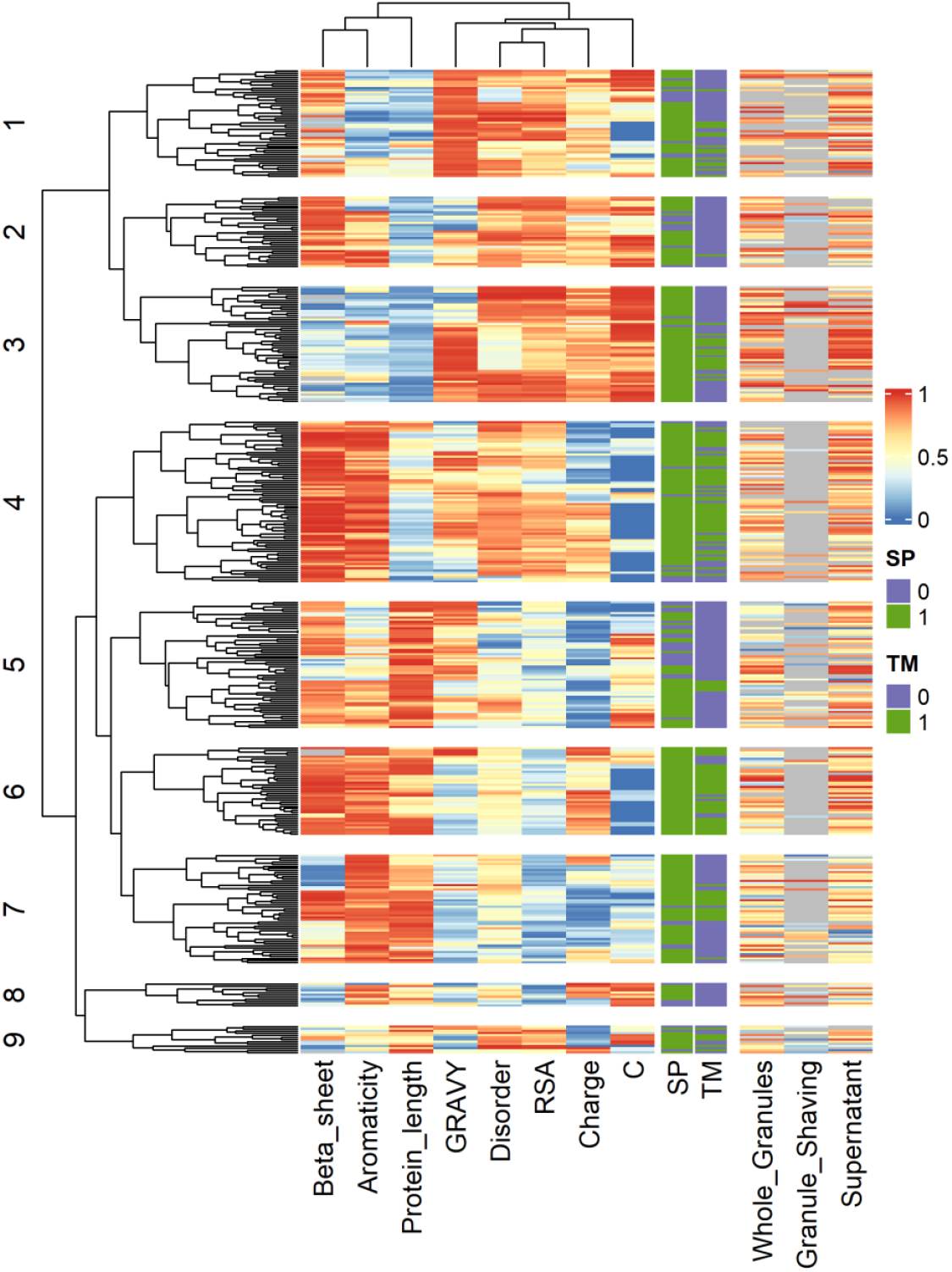
Clustering of proteins according to aggregating properties. Clustering of the different protein groups (y-axis) using the ranked sequence-based properties (x-axis; colours). The proteins were classified based on the following aggregate forming properties: the beta-sheet percentage, aromaticity, protein length, hydrophobicity (GRAVY), disorder, relative surface accessibility (RSA), net charge, and cysteine percentage. The protein groups were annotated based on the presence of a signal peptide (SP), a transmembrane domain (TM) and the ranked protein abundance for the different approaches. Detailed information about each cluster can be found in Supplemental Table 2.

Cluster 1 and 2 mainly contained short proteins with high surface accessibility. In Cluster 1, the proteins were also hydrophobic, and the most abundant protein contained a PEP-CTERM motif and clustered with proteins with similar motifs. The proteins in Cluster 2 additionally had a high beta-sheet percentage. The most abundant protein in Cluster 2 contained a Phage_T4_gp19 domain and clustered together with proteins containing a T6SS_HCP domain, which suggests that these proteins are subunits for different tubular proteins (IPR010667; IPR008514). Similar to the first clusters, Cluster 3 contained short proteins, but they had a low beta-sheet percentage and a high cysteine percentage. The most abundant proteins in this cluster were pilus and cytochrome C proteins. The proteins in Cluster 4 were rather the opposite of proteins in Cluster 3 as they were generally longer, had a low cysteine percentage, a high beta-sheet percentage and a high aromaticity. The proteins in this cluster were mainly beta-barrel membrane proteins, including porins and outer membrane proteins. Interestingly, this cluster also contained a Flavobacterium protein with a curlin repeat domain which is involved in cell adhesion (IPR009742). The proteins in Cluster 6 had similar properties as those in Cluster 4 and also contained abundant beta-barrel membrane proteins, although these proteins were longer. The properties in Cluster 7 resembled those in Cluster 6, except that the proteins in Cluster 7 had a lower beta-sheet percentage and a negative charge. The most abundant proteins in this cluster bind extracellular solutes, including starch, or have a sulfatase domain. The proteins in Cluster 5 were long and negatively charged, but relatively hydrophobic and contained a relatively large fraction of beta-sheets. The two most abundant proteins in this cluster were labelled hypothetical but showed decent sequence homology (49–58%) against proteins from the Paracidovorax genus labelled as cell surface proteins. Furthermore, this cluster contained multiple proteins with a polycystic kidney disease (PKD) domain, which are mainly found in large extracellular cell surface glycoproteins (IPR000601), multiple proteins with a fibronectin domain that is among others involved in protein binding and cell adhesion (IPR003961), and multiple proteins annotated as outer cell wall protein that shared sequence homology (32–33%) with proteins from the Bacillota phylum containing an S-layer homology domain. Remarkably, despite the high enrichment of Accumulibacter (75–80%), most proteins in this cluster did not originate from Accumulibacter (Supplemental Table 2). Clusters 8 and 9 were the smallest clusters and mainly contained proteins with a high positive or negative charge, with Cluster 8 containing proteins with a low surface accessibility without transmembrane domain and Cluster 9 containing proteins with a high surface accessibility. The most abundant proteins in these clusters did not have similar functions, indicating that these clusters were not yet well defined.

In total, our identified proteins covered between 55 and 70% of the secreted and extracellular protein biomass. Clusters 3 and 6 were the most abundant in the supernatant and whole granules, although Cluster 6 was over 2-fold more abundant than Cluster 3 in whole granules (Supplemental Figure 6). Conversely, only Cluster 3 was highly abundant in shaved granules. Furthermore, Cluster 1 was more abundant in the supernatant compared to the whole and shaved granules. Taken together, we identified different clusters of proteins based on physicochemical properties, which included abundant proteins such as pili and porins.

## DISCUSSION

We employed two advanced approaches that successfully enriched for extracellular and secreted proteins when compared to the whole granule approach. This obtained enrichment also exceeded the enrichment observed when extracting extracellular polymeric substances in alkaline conditions at high temperatures [27, 28]. Nevertheless, a substantial number of intracellular proteins was identified by both approaches that were employed in this study, although their low abundance indicates that these proteins are likely released in the extracellular space following cell lysis [53].

Interestingly, both approaches enriched for different proteins. The enzymatic granule shaving approach uniquely identified multiple cytochrome C proteins, but could not observe transmembrane and structural proteins, which are challenging to digest in their native form. We found that post-shaving digestion under denaturing conditions (6 M urea) tremendously increases the reproducibility and number of identified peptides. As most proteases are not resistant to high urea concentrations, other denaturing agents such as Rapigest might improve the identification of these proteins during the shaving treatment [54]. In the supernatant, the fraction of Accumulibacter proteins was smaller compared to the granule-based samples. The lower Accumulibacter enrichment could be the result of non-Accumulibacter planktonic cells present in the reactor, even though we aimed to remove them by centrifugation. Therefore, it might be more effective to select against these cells in the culture themselves, for example by shorter settling or shortening the aerobic phase. Alternatively, the low Accumulibacter enrichment might originate from the taxonomic classification method. We have identified multiple high abundant proteins that were not assigned to Accumulibacter, even though their sequence corresponded to Accumulibacter. The taxonomic classification might improve when using a less strict least common ancestor approach or when performing taxonomic classification on the contigs or even metagenome-assembled genome level [11, 12]. This would result in more clarity about the genes originating from a single organism.

Using the advanced approaches, we aimed to identify potential aggregate forming proteins that could play a structural role in the biofilm matrix. Various biofilm components have been identified before to play a role in biofilm formation, stabilization and structure [1, 2, 14] and our approaches have identified various proteinaceous components. For example, we identified filamentous phage tails and other tube-forming proteins. Such proteins have been previously identified in *Pseudomonas aeruginosa* biofilms to promote organised and stable biofilms [55]. In addition, we identified pili proteins which are reported play a role in bacterial self-organisation in biofilms and stability of the biofilm [56]. In that same cluster, we found cytochrome C proteins, which may be stacked by their heme groups thereby forming pili-like structures called nanowires [57, 58]. The location of cytochrome C could be further examined by staining and visualization. Interestingly, phage tail, pili, and cytochrome C proteins have been reported to be more expressed in granular compared to floccular Accumulibacter-enriched cultures, suggesting that these filamentous structures indeed play a role in granule formation [26].

In addition to filamentous proteins, we identified multiple clusters that contained porins and other transmembrane beta-barrel proteins, which form membrane channels that allow the diffusion of small molecules. However, rising evidence indicates that these beta-barrel proteins can form functional amyloids [59-62], which play pivotal roles in biofilm formation and stabilization [14, 63]. Our dataset might shine some light on this hypothesis as our supernatant approach is designed to enrich for non-transmembrane proteins over transmembrane proteins, suggesting that the identified beta-barrels could be functional amyloids. Interestingly, we indeed identified an amyloid curli protein among these beta-barrel proteins in the supernatant but not in whole granules. This curli protein originated from Flavobacterium, a genus that was 1.7-fold more present in the supernatant compared to the granules, which might explain why it was not identified in the whole granules. To further investigate the presence of amyloid beta-barrels in the supernatant, it would be interesting to stain the supernatant proteins for amyloid structures before and after digestion to correct for other amyloid structures that are resistant to denaturation and proteases and therefore not identified [13].

We also found proteins that have been previously associated with biofilm formation, although their function is less clear. For example, we identified proteins with PEP-CTERM motifs that are exclusively found in bacteria that have both an outer membrane and exopolysaccharide production genes [64]. Moreover, these proteins have been reported to play a role in floc formation in other bacteria found in wastewater treatment plants [65-67] and have been found overexpressed in granular compared to floccular Accumulibacter-enriched cultures [26]. Therefore, further unravelling the function of this motif may advance our understanding of the granule formation. In addition, we identified proteins with polycystic kidney disease (PKD) and fibronectin 3 (fn3) domains which clustered together with proteins homologous to proteins with an S-layer domain. While all these proteins are linked to the cell surface [68], S-layer proteins have been found in other places in the biofilm, aiding in bacterial assembly and aggregation [69] and host fibronectin was reported to promote aggregation in *Staphylococcus aureus* [70, 71]. This might suggest that proteins with PKD and fn3 domains could also play a role in biofilm formation beyond cell adhesion.

For most clusters, at least one of the highly abundant proteins was attributed to Accumulibacter, confirming that Accumulibacter is a main contributor to the extracellular matrix of the granules. However, the clusters with shorter beta-barrels (Cluster 4) and cell surface proteins (Cluster 5) mainly contained proteins from other organisms. This suggests that proteins from multiple organisms may influence the granule formation and structure, where Accumulibacter mainly contributes various types of filamentous proteins and other community members contribute cell surface proteins and potentially amyloid-like proteins. Since the formation of dense granules is linked to Accumulibacter enrichment [24] and to an increased filamentous protein production [26], filamentous protein production might be an important player in granule formation. The aggregating role of filamentous proteins like pili has been described in the formation of microcolonies [56], but the complex and interactive nature of biopolymers in the biofilm matrix complicates the characterisation of single protein types.

In conclusion, the two advanced metaproteome approaches successfully enriched the extracellular metaproteome from a *Ca*. Accumulibacter enrichment, which is crucial for understanding biofilm and granule formation, as well as for optimizing or even controlling microbial communities in medical and industrial settings. Among many yet uncharacterised proteins we also identified several proteins with properties that were previously linked to granule or biofilm formation and biofilm structure. This data resource is a starting point to further unravel the influence of proteinaceous components on biofilm and granule formation.

## Supporting information

SI DOC

## ACKNOWLEDGEMENTS

The authors acknowledge Dita Heikens for her support with sample preparation, Ramon van der Zwaan for insightful discussions about metaproteomics, and all other colleagues from the Department of Biotechnology for valuable discussions. This work was supported by Novo Nordisk Foundation (grant NNF22OC0071498). Figure 1A was created in BioRender (https://BioRender.com/h86r472).

