## Supplementary material for "Metaproteomic profiling of the secretome of a granule-forming *Ca*. Accumulibacter enrichment": SI DOC

#### Table of contents

| | | |
|---|---|---|
| **Supplemental Figure 1** | Sample reproducibility for whole granule lysates | 2 |
| **Supplemental Figure 2** | The influence of different proteases treatments on granule shaving | 3 |
| **Supplemental Figure 3** | The influence of sampling phase for the supernatant approach | 5 |
| **Supplemental Figure 4** | Global comparison of different metaproteomics approaches | 6 |
| **Supplemental Figure 5** | Enrichment of proteins at different cellular locations | 7 |
| **Supplemental Figure 6** | Distribution of the structural biomass over the different clusters | 8 |

### Sample reproducibility for whole granule lysates

The reproducibility of the whole granule samples was examined as a reference for the other two approaches and was tested on both a protein extraction and on the MS measurement level (multiple injections of the same sample). When injecting the same sample digested with trypsin twice, up to 80% of the protein groups were identified in both replicates (Supplemental Figure 1A) and the abundances correlated well (average Pearson: 0.95; Supplemental Figure 1B). The protein groups identified in a single replicate were low abundant. Extracting two samples from the same reactor resulted in a similar reproducibility with 77% of the protein groups being identified in both replicates and the abundances correlating well (average Pearson 0.93). Over all four samples, 2767 protein groups were identified of which 60% was identified in all samples with good correlation (average Pearson: 0.94) and 15% of the protein groups was found in single replicates. Digesting the samples with chymotrypsin instead of trypsin resulted in 1085 protein groups of which only 66% was identified in both replicates (Supplemental Figure 1A), albeit at good reproducibility (average Pearson: 0.90). Interestingly, the chymotrypsin samples identified 52 protein groups that were not identified in the trypsin samples, resulting in 2819 proteins identified in whole granules. Taken together, our method for analysing the proteome from whole granules is reproducible, especially when digesting with trypsin.

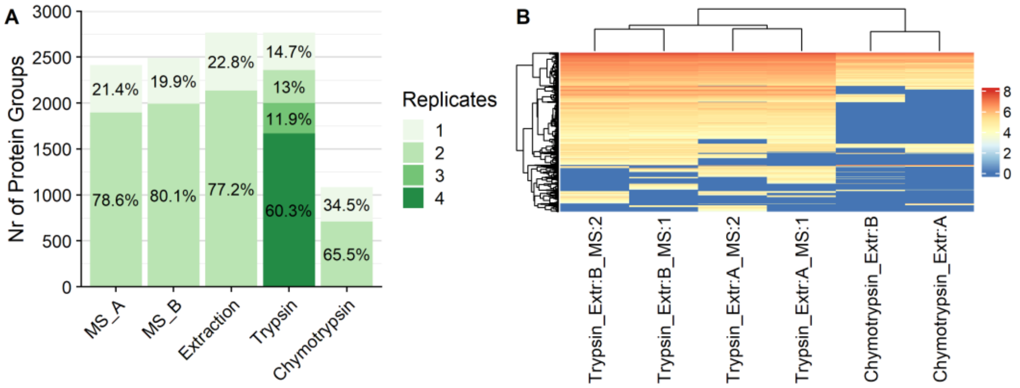


***Supplemental Figure 1: Sample reproducibility for whole granule lysates.*** *(A) Number of identified protein groups (y-axis) shared between replicates (colours) when injecting the same sample twice (x-axis; "MS"), when extracting two samples from the same reactor (x-axis; "Extraction") and when combining all samples digested with trypsin (x-axis; "Trypsin") or chymotrypsin (x-axis; "Chymotrypsin"). (B) Correspondence of the different samples as indicated by hierarchical clustering. The x-axis indicates the protease, the extracted sample [Extr:A or Extr:B] and the MS injection [MS:1 or MS:2]*

### The influence of different proteases treatments on granule shaving

The granule shaving approach relies on the digestion of native proteins, which are not easily accessible for proteases. Therefore, the identification of native proteins was enhanced by shaving the granules with different proteases in parallel. In addition, more miscleavages were allowed, but only a limited number of peptides contained more than two missed cleavages (Supplemental Figure 2C). Shaving the granules with trypsin identified 414 protein groups (Supplemental Figure 2A), whereas initial shaving with LysC followed by a combined shaving of LysC and trypsin resulted in 543 protein groups, of which 285 overlapped between the protease treatments with a moderate correlation (average Pearson: 0.66). Nevertheless, almost half of the identified protein groups shaved with LysC+Trp was only identified in single replicates (not the same replicate; Supplemental Figure 2B), making shaving with trypsin slightly more reproducible (35% in single replicates), although the reproducibility was more variable than whole granule samples. Shaving the granules with chymotrypsin yielded only 91 protein groups of which 75% was identified in only one replicate, which is the most variable result in the shaving approach. This is most likely explained by the self-digestion of chymotrypsin due to the low protein availability and the elevated digestion temperature. Nevertheless, each protease treatment identified unique protein groups that were not identified by the other proteases, highlighting the benefit of combining the different protease treatments. However, all these treatments displayed a large amount of undigested protein at high retention times (Supplemental Figure 2D). Therefore, the identification of native proteins was further enhanced by exposing proteins that were solely shaved with LysC to denaturing conditions (6M Urea) during the continued digestion after granule removal. The denaturing conditions indeed resolved the large amount of undigested protein (Supplemental Figure 2D). Furthermore, this treatment identified 655 protein groups of which 68% was found in at least two replicates, making it the shaving treatment with the most identifications and the highest reproducibility. Moreover, almost half of the identified protein groups was not identified by the other proteases treatments, underpinning the added benefit of post-shaving denaturation in identifying proteins by shaving. Overall, the processing of native proteins makes the shaving approach less reproducible than the WG approach. However, their identification is greatly enhanced by using multiple proteases in parallel, identifying in total 973 protein groups.

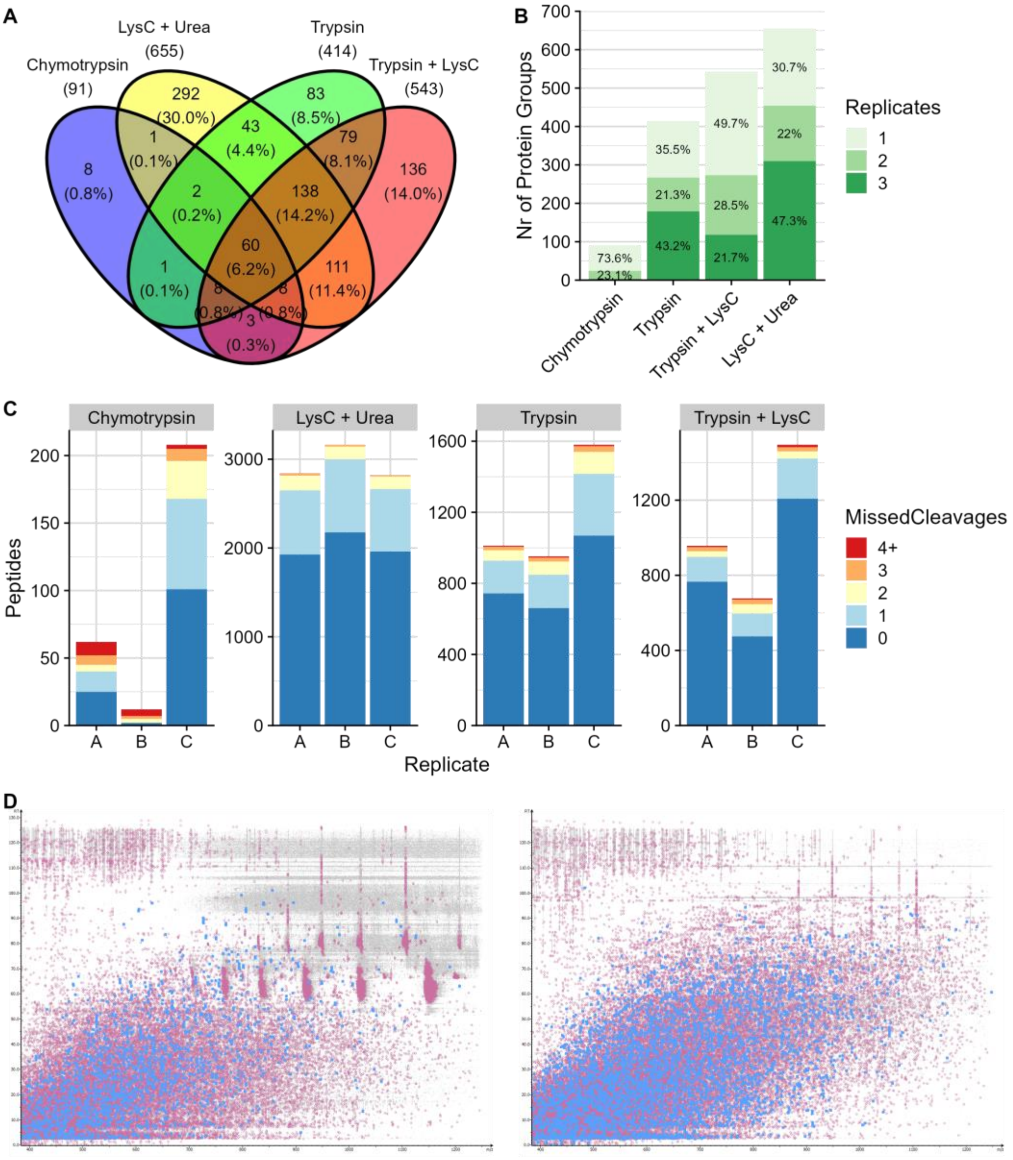


***Supplemental Figure 2: The influence of different protease treatments on granule shaving.*** *(A) Venn diagram displaying the identified protein groups per protease treatment and the overlap between them. (B) Number of identified protein groups (y-axis) shared between the replicates (colours) of the different protease treatments (x-axis). (C) Number of peptides (y-axis) with a specific number of missed cleavages (colours) for the different replicates (x-axis) per treatment (panels). (D) Retention time (y-axis) against mass over charge (x-axis) displaying the features (purple) and identified peptides (blue) of representative samples of the Trypsin+LysC treatment (left panel) and the LysC+Urea treatment (right panel). The left panel shows a large amount of indigested protein which are not present in the right panel.*

### The influence of sampling phase for the supernatant approach

The supernatant approach identifies proteins in the culture's supernatant, whose protein content is expected to be most concentrated at the end aerobic growing phase, right before discharge of the supernatant and refill with new medium. However, the protein content might differ between the anaerobic feeding phase and after the aerobic growing phase. Therefore, we identified the proteins in the supernatant obtained after each phase. Most protein groups (1398) were found in both phases with good correlation (average Pearson: 0.86; Supplemental Figure 3A), although 156 and 485 protein groups were uniquely found in the anaerobic and aerobic phase, respectively. Furthermore, the protein identification was slightly more reproducible, and the protein biomass was 1.6-fold higher in the aerobic phase compared to the anaerobic phase (Supplemental Figure 3B). However, these additional protein groups in the aerobic phase could represent intracellular proteins, while we are interested in secreted proteins. To distinguish the extracellular from the intracellular proteins, we predicted the presence of a transmembrane domain and a signal peptide for secretion based on the protein sequence. The abundance of these proteins relative to the protein biomass was similar in both supernatant types (Supplemental Figure 3C), indicating that both types identify a similar number of secreted proteins. Taken together, the supernatant at the end of the aerobic growing phase results in more proteins identifications, higher reproducibility, and a similar number of extracellular proteins compared to the anaerobic phase. However, as there are some proteins uniquely identified in the anaerobic phase, we regarded the samples from both phases as replicates in further analyses.

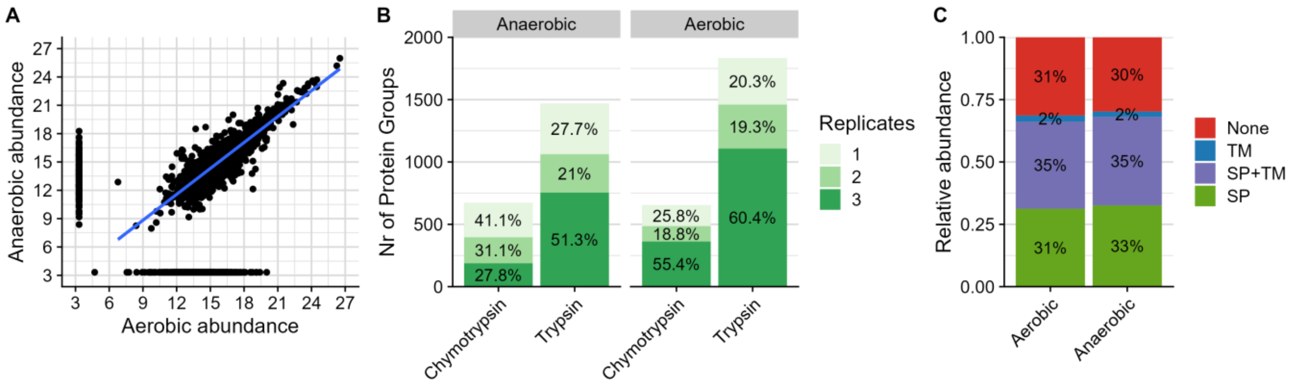


***Supplemental Figure 3: Influence of sampling phase on the supernatant metaproteome.*** *(A) Correlation between aerobic (x-axis) and anaerobic (y-axis) samples. (B) Number of identified protein groups (y-axis) shared between the replicates (colours) for both sampling phases (panels) and digestion treatments (x-axis). (C) Enrichment of the relative protein abundance for secreted proteins based on the presence of a signal peptide (SP) and/or transmembrane domain (TM; colours) for both sampling phases.*

### Supplemental Figures

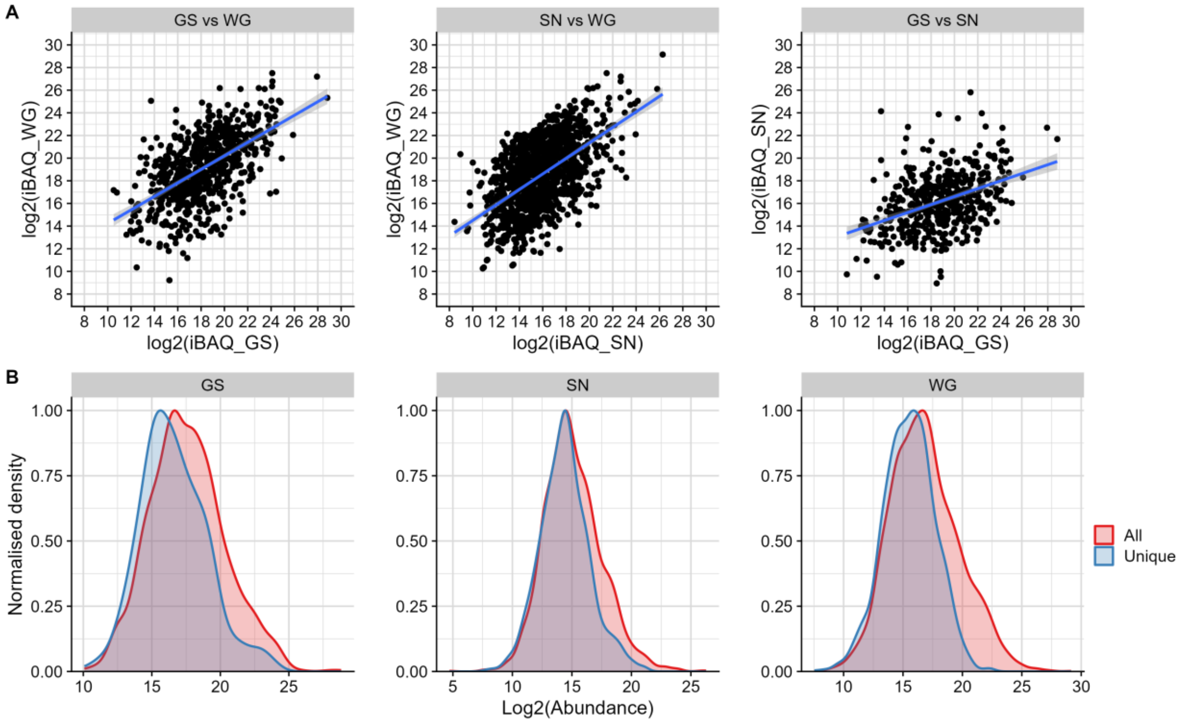


***Supplemental Figure 4: Global comparison of different metaproteomics approaches.*** *(A) Correlation between the log2 protein abundances from the different approaches (B) Overlap in the log2 protein abundance (x-axis) for all protein groups (red) and unique protein groups (blue) identified by the different approaches (panels).*

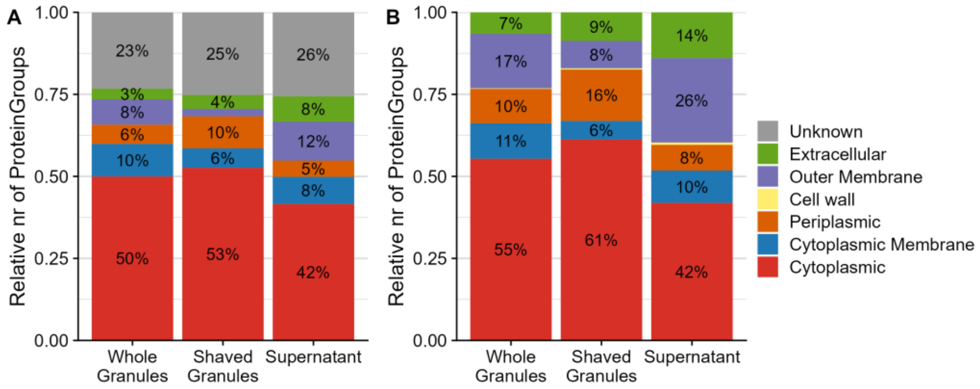


***Supplemental Figure 5: Enrichment of proteins at different cellular locations.*** *The relative number of protein groups (y-axis) predicted at different cellular locations (colours) as predicted by (A) PSORTb and (B) DeepLocPro for the different approaches (x-axis).*

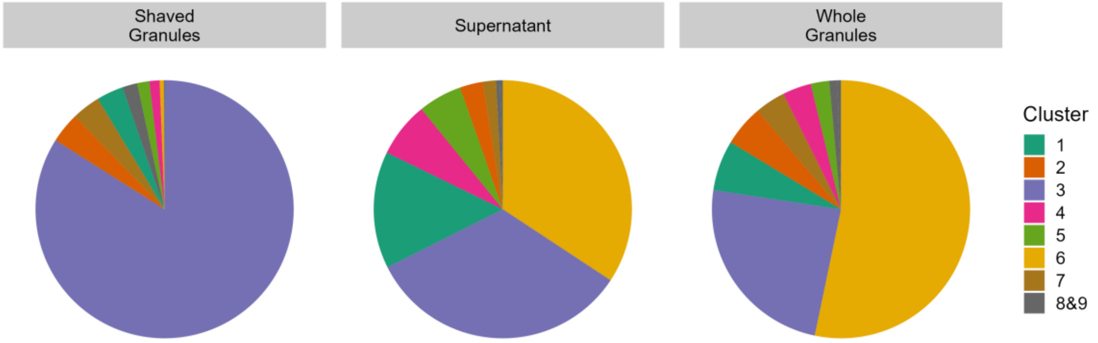


***Supplemental Figure 6: Distribution of the structural biomass over the different clusters.*** *The abundance of the different clusters (colours) relative to the total structural biomass for the different approaches (panels).*
